# Cell Mechanics at the Rear Act To Steer the Direction of Cell Migration

**DOI:** 10.1101/443408

**Authors:** Greg M. Allen, Kun Chun Lee, Erin L. Barnhart, Mark A. Tsuchida, Cyrus A. Wilson, Edgar Gutierrez, Alexander Groisman, Alex Mogilnerd, Julie A. Theriot

**Affiliations:** Department of Biochemistry and HHMI, Stanford University School of Medicine, Stanford, CA 94305 USA; Department of Neurobiology, Physiology and Behavior, University of California, Davis, CA 95616 USA; Department of Physics, University of California, San Diego, CA 92023 USA; Courant Institute of Mathematical Sciences and Department of Biology, New York University, New York, NY 10012 USA; Department of Biology and HHMI, University of Washington, Seattle, WA 98105 USA

**Keywords:** self-organization, cell turning, cell migration, cell motility, keratocyte, actin, myosin, adhesion, asymmetry

## Abstract

Motile cells navigate complex environments by changing their direction of travel, generating left-right asymmetries in their mechanical subsystems to physically turn. Currently little is known about how external directional cues are propagated along the length scale of the whole cell and integrated with its force-generating apparatus to steer migration mechanically. We examine the mechanics of spontaneous cell turning in fish epidermal keratocytes and find that the mechanical asymmetries responsible for turning behavior predominate at the rear of the cell, where there is asymmetric centripetal actin flow. Using experimental perturbations we identify two linked feedback loops connecting myosin II contractility, adhesion strength and actin network flow in turning cells that are sufficient to recreate observed cell shapes and trajectories in a computational model. Surprisingly, asymmetries in actin polymerization at the cell leading edge play only a minor role in the mechanics of cell turning – that is, cells steer from the rear.

**Highlights:**

- Fish keratocytes can migrate with persistent angular velocity, straight or in circles.
- Asymmetry in protrusion at the leading edge is not sufficient to generate persistent turning.
- Asymmetries in myosin II contraction, actin flow and adhesion at the cell rear cause turns.
- Our new computational model of migration predicts observed cell trajectories.

## Introduction

Directed cell migration is required for many fundamental processes in multicellular animals, including wound healing, immune cell trafficking and embryonic development. In order to achieve these complex behaviors, cells must have both an ability to translocate in space and a mechanism to determine and alter the direction of this migration. Previous work has established a good understanding of the physical mechanisms that drive steady state migration (Blanchoin et al., 2014; Lauffenburger and Horwitz, 1996; Mogilner and Rubinstein, 2010; Pollard and Borisy, 2003) while also mapping out the signaling pathways that dictate the choice of direction of motion in response to environmental cues (Jin, 2013; Parent and Devreotes, 1999; Ridley et al., 2003; Schneider and Haugh, 2006; Xiong et al., 2010). However, the physical mechanisms that cells use to transduce asymmetries in signaling pathways into the coordinated large-scale reorientation of the entire motile cell remain obscure. In this work we seek to map out how an individual cell physically adapts the machinery of cell motility to change its direction of travel while maintaining persistent polarization.

All migrating cells are structurally polarized along the direction they migrate. Actin-based locomotion typically involves preferential localization of the assembly of actin filaments to the front of the cell (Theriot and Mitchison, 1991; Wang, 1985) while the cell body and the contractile activity of non-muscle myosin-II typically localize to the rear (Vicente-Manzanares et al., 2007; Wilson et al., 2010; Yumura et al., 1984). This polarization is sufficient to weakly determine the direction that a cell will travel at a given instant in time (Jiang et al., 2005); yet, over a distance of even a few cell lengths (~100 *μ*m) no cell follows a completely persistent path in space (Arrieumerlou and Meyer, 2005; Dunn, 1983; Li et al., 2008). In principle, transient left-right asymmetries in any of the mechanical components contributing to directed migration including protrusion at the leading edge, retrograde flow of the polymerizing actin network, adhesive coupling to the substrate, retraction at the rear, or contraction of the cell body could cause slight changes in the overall orientation of a migrating cell and alter its direction of movement. If these variations are random and uncorrelated, as might be expected for a cell moving spontaneously in a uniform environment, the resultant path of motion would be a persistent (or correlated) random walk (Dunn, 1983). Indeed, persistent random walk statistics are approximately able to fit the long-distance trajectories observed for many types of cells moving spontaneously in two dimensions, although detailed analysis often reveals that there must also be non-random components (Hartman et al., 1994; Selmeczi et al., 2005; Stokes et al., 1991)

Migrating cells can also actively change direction in response to spatial cues from their environments, including signals from soluble or matrix-attached chemical gradients, or even electric fields (Cortese et al., 2014; Ridley et al., 2003). For rapidly moving cells that tend to maintain strong front-rear polarization, such as human neutrophils and fish epidermal keratocytes, motile cells exposed to a rapid change in their primary directional cue tend to reorient their migration in a gradual U-turn rather than depolarizing and then repolarizing along a new front-rear axis (Cooper and Schliwa, 1986; Xu et al., 2003; Zigmond et al., 1981). This observation indicates that rapidly motile, persistently polarized cells are capable of generating persistent mechanical left-right asymmetries that result in cell turning.

Most studies of the mechanics of cell turning during persistent migration have focused on characterization of asymmetries at the cell leading edge (Insall, 2010). For example, the direction of travel in fibroblasts and endothelial cells is thought to be altered by the summation of competing weakly stable PI3K dependent protrusive branching events (Weiger et al., 2010; Welf et al., 2012), with a relationship between asymmetric calcium flickers and the direction of travel (Tsai and Meyer, 2012; Wei et al., 2009). Work in axonal growth cones of neurons has also implicated local variation in microtubule dynamics (Buck and Zheng, 2002) and adhesion turnover (Myers and Gomez, 2011) in turning behavior. This prior work examining the leading edge, though, has not taken into account the complete set of physical interactions inherent to turning in cell migration, as significant mechanochemical systems that orchestrate cell motility exist outside of this region. For instance it has been established that the initial process of polarization can originate at the cell rear (Cramer, 2010; Mseka et al., 2007; Yam et al., 2007), and that positive feedback between myosin contraction and actin flow and negative feedback between actin flow and adhesion are responsible for this polarization (Barnhart et al., 2015). Furthermore, it has not been possible to use the mechanics of cell migration to recreate the detailed trajectories that cells have been observed to take when only the behavior of the leading edge is taken into account (Selmeczi et al., 2005; Stokes et al., 1991).

In order to develop an understanding of the cell-scale mechanics necessary to change the direction of travel, we have used the model system of the fish epidermal keratocyte. These rapidly-moving cells are mechanically and geometrically simpler than fibroblasts or neutrophils, allowing construction of quantitative mechanistic models that can replicate even complex aspects of their observed behavior (Barnhart et al., 2011; Keren et al., 2008). Motile keratocytes also normally maintain only a single protrusion with an extremely long persistence time, making them an ideal system to study the methods for mechanical determination of left-right turning behavior at steady state.

Migration in keratocytes is governed by a set of fundamental forces and mechanical actors that have been integrated into a general mechanical model (Barnhart et al., 2011). Driving protrusion is a densely branched polymerizing dendritic actin network at the leading edge (Svitkina et al., 1997). Opposing the force of actin polymerization is the tension in the plasma membrane of the cell, which spatially integrates force across the cell (Keren et al., 2008). Nascent adhesions are laid down at the leading edge (Lee and Jacobson, 1997) and coordinate the protrusive force of the actin network into mechanical work with only a minimal rate of slip, otherwise known as retrograde flow (Wilson et al., 2010). Adhesions chemically mature and become mechanically weaker as the cell translocates over them (Barnhart et al., 2011). A large set of binding proteins interact with the actin network to bundle filaments, cap barbed ends, create new branch points, and sever filaments. This set includes the force-generating protein myosin II, which binds the actin network and acts to contract and disassemble it at the rear of the cell (Wilson et al., 2010). This contraction of the actin network creates retrograde flow of filamentous actin at the cell rear oriented towards the cell body and perpendicular to the direction of motion.

Prior examination of keratocyte turning has shown a correlation between asymmetric traction stresses and sharp turns (Oliver et al., 1999) as well as an asymmetry in the coupling of actin filament motion and traction forces (Fournier et al., 2010). However, these prior analyses were limited in scope to only the forces applied to the substrate. In this work we seek to determine how all known mechanical actors of cell motility (actin network assembly, network disassembly, network contraction, adhesion to the substrate and membrane tension) produce lateral asymmetry and consequently turning behavior in motile keratocytes. Surprisingly, we find that left-right asymmetries determining turning behavior are controlled by the interwoven actions of myosin II contraction and substrate adhesion at the rear of the cell, and not by actin polymerization at the front of the cell, producing a form of “rear-wheel steering”. These mechanical systems are organized in a combination of feedback loops, such that these cells can enter into and exit from states of stable persistent turning.

## Results

### Keratocytes enter into persistent turning states

To begin to explore the physical mechanisms that underlie cell turning, we first measured the trajectories of 38 spontaneously migrating individual cells over time periods of four to eighteen hours (**Figure 1A**). While a minority of cells (9 of 38 examined) seemed to follow a meandering trajectory approximating a persistent random walk, the majority (29 of 38) exhibited trajectories that included at least some strikingly persistent segments, where the cell typically turned in a small circle several times, intermixed with periods of nearly straight movement, reminiscent of “knots on a string” (**Figure 1B**). These two qualitatively distinct types of motion could be distinguished quantitatively by comparing their angular speed (*ω*) over time and the autocorrelation of the angular speed *A*(*ω*) (**Figure 1C-E**). Persistent cells (e.g. cell-7; magenta) typically moved with a fairly constant angular speed for many minutes before switching (**Figure 1C**), generating strong positive autocorrelation signals (**Figure 1D**) and bimodal distributions of angular speeds (**Figure 1E**), while non-persistent cells (e.g. cell-8; gold) showed a much faster decay in autocorrelation and a unimodal distribution of angular speeds centered around 0. Persistently turning cells were equally likely to travel in clockwise or counter-clockwise circles, and individual cells could switch between CW and CCW turns without apparent bias or memory (**Figure S1A**). The dwell time between switching events was broadly distributed (**Figure S1B**).

**Figure 1.**
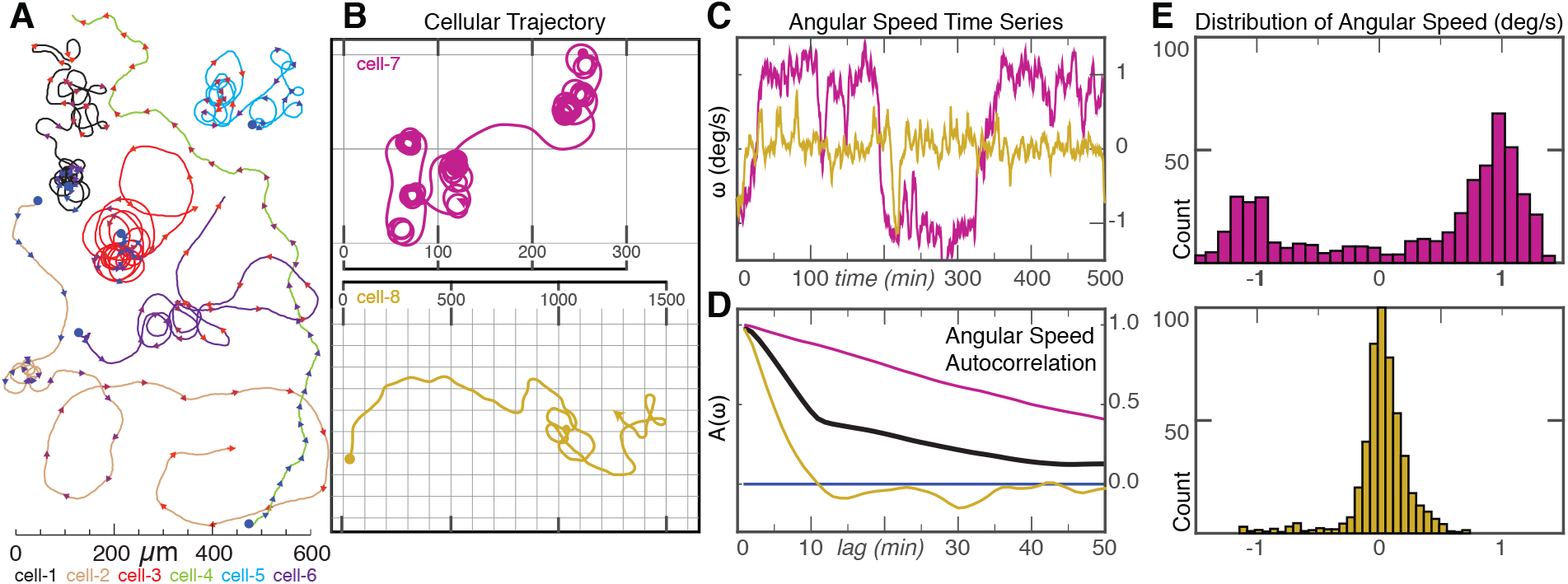
Long-term trajectories of single cells exhibit persistent turning states. **(A)** Example trajectories of 6 keratocytes cells over a time scale of ~10 hours. Trajectories start at blue dots, and each color represents a different cell. Arrowheads indicate the current direction of motion every 20 minutes and are colored from blue/start to red/finish. Scale bar indicates distance traveled. Note cells can exhibit two phases of migration with long periods of persistent turning intermixed with periods of straighter paths. **(B)** Example trajectory of a cell exhibiting prolonged periods of persistent turning (*cell-7, magenta, top*) and a cell following a wandering path (*cell-8, gold, bottom*). Scale in micrometers. **(C)** Time series of angular velocity, (*ω*), for cell-7 (*magenta)* and cell-8 (*gold)*. **(D)** Auto-correlation of angular velocity (*A*(*ω*)) as a function of lag time in minutes for cell-7 (*magenta)*, cell-8 (*gold)* and average of all 38 cells (*black*). **(E)** Distribution of angular velocities for the persistently turning cell-7 (*magenta, top*) and the wandering cell-8 (*gold, bottom*).

Using simulations of randomly generated trajectories, we explored whether any kind of simple stochastic fluctuations could account for these observed behaviors (**Figure S1C**). A simple random walk model where the direction of motion is randomized at each time step (**Figure S1C, Model A**) gives a uniform distribution of angular speeds, and no persistent directional autocorrelation. A more realistic model of a persistent random walk in which a cell tends to continue moving in the same direction but with slight random turns at each step, such as has been used to describe the motility of *Dictyostelium discoideum* (Van Haastert, 2010), fibroblasts (Gail and Boone, 1970), endothelial cells (Stokes et al., 1991) and granulocytes (Hall and Peterson, 1979) (**Figure S1C, Model B**) could produce trajectories that resembled those of the subset of less persistent keratocytes with an appropriate distribution of angular speeds, but with less angular autocorrelation than was experimentally observed. In an attempt to appropriately simulate the trajectories of the majority of keratocytes with persistent turning and high *A*(*ω*), we next attempted simulating stochastic variations in the time derivative of *ω* rather than in *ω* itself (**Figure S1C, Model C**). This model, in essence a correlated random walk in angular velocity space, has been used previously to describe the paths of humans who have been blindfolded to their external environment, who frequently generate trajectories qualitatively similar to “knots on a string” (Souman et al., 2009), an interesting parallel to the non-chemotactic keratocytes. The blindfolded human model produced trajectories that were somewhat reminiscent of the keratocytes and did exhibit slow autocorrelation decay of angular speed, but could not reproduce the frequently observed bimodal distribution of instantaneous angular speeds (**Figure S1C, Model C**). The nature of the quantitative mismatch between this model and the experimental observations can also be seen by examining the mean squared displacement of angular speed over time; the model shows a diffusive (approximately linear) relationship while the experimental data for keratocytes quickly rises and then plateaus (**Figure S1D**). Thus, the observed keratocyte behavior cannot be adequately explained by any simple fluctuation-based model.

Mechanically, the existence of persistently turning keratoctyes with bimodally distributed angular speeds informs us that there must exist persistent asymmetries in mechanical behavior that are perhaps maintained by feedback of the act of turning on the mechanical asymmetries that produce turning. Individual cells with particularly strong autocorrelations in angular speed were also typically fast-moving and typically turned in tight circles (**Figure S1E,F**), although in the population as a whole there was no strong correlation between net speed and angular velocity (**Figure S1G**).

### Turning cells exhibit an asymmetric shape and F-actin distribution

The observation that many keratocytes are able to enter into a persistently turning state afforded us the opportunity to examine cytoskeletal variations associated with cell turning. In order for cells to enter into persistent turns, there must be a persistent imbalance in at least one of the mechanical elements that produce motility. Images of a typical persistently turning cell (**Figure 2A,B and Movie S1**) showed that the cell body at the rear is centered toward the inner side of a turn, in contrast to the typical symmetric fan shape of a keratocyte that is moving in a straight line (Csucs et al., 2007; Goodrich, 1924). The flat, actin-filament rich lamellipodium at the front of the turning cell is also not symmetric, but instead is elongated in the “wing” on the outer side. Whole cells with an overall increased aspect ratio (elongated lamellipodia) tend to also exhibit increased cellular speed (Keren et al., 2008) suggesting that there may be asymmetric protrusion along the leading edge for turning cells, such that the part of the lamellipodium on the outside of the turn is effectively moving faster than the part on the inside.

**Figure 2.**
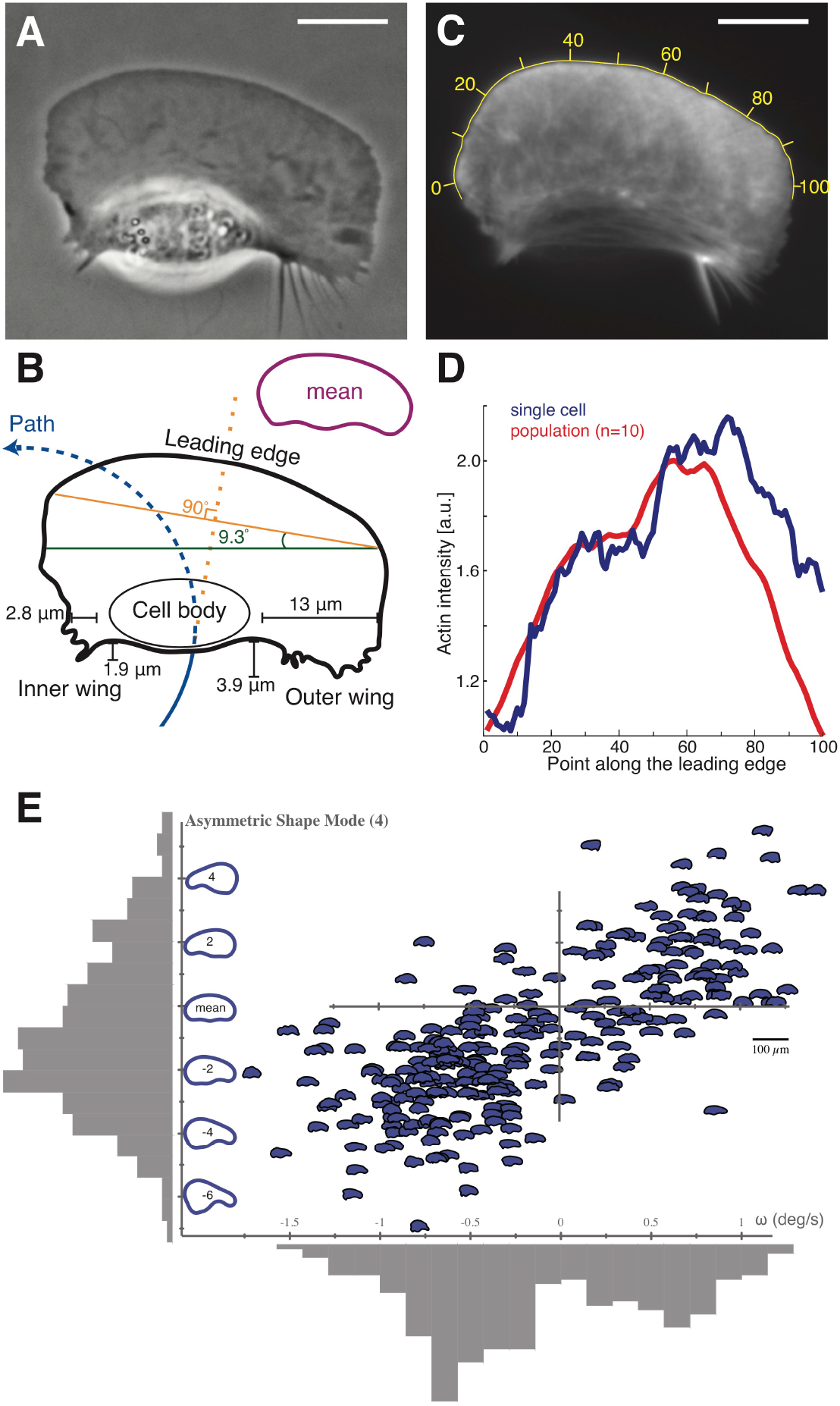
Turning cells have asymmetric shapes and F-actin distributions. **(A)** Example phase-contrast image of a cell in an asymmetric circular path turning counter-clockwise. Scale bars in all panels are 10 *μ*m. **(B)** The contour of this cell illustrating the cell path (*blue line*) as well as the elongated aspect ratio on the outer side of the turning cell. The cell body is displaced towards the inside of the turn, and the outer wing lags behind the cell. The leading edge orientation (*orange line*) is orthogonal to the direction that the cell was previously traveling previously (*dashed orange line*) and the rear edge orientation (*green line*) is orthogonal to the direction the cell is currently traveling. The inset contour shows the average shape of 22 mutually aligned turning cells. **(C)** F-actin distribution from this turning cell as visualized by AF-488 phalloidin labeling, with yellow numbers indicating position along the leading edge. **(D)** Measured density of F-actin along the points of the leading edge of the cell in panel C (*blue line*) and in the average of a population of 10 turning cells (*red line*). Note the asymmetric accumulation of actin filaments at the leading edge on the outer side. **(E)** For a single cell that is being forced to turn at a high rate by exposure to multiple external electric fields over 272 time points over approximately 20 minutes, the angular velocity at each time point, *ω*, is plotted against the left-right asymmetric PCA shape mode (Keren et al., 2008) as depicted on the vertical axis. The distribution of values for both asymmetric shape and angular velocity are plotted in grey adjacent to each axis. Calculated correlation coefficient is 0.73, calibration bar for cell outline images is 100 *μ*m.

The shape of the turning cells suggests that there are two relevant axes that describe the shape of the cell relative to its trajectory. One is the orientation of the leading edge, as defined by the front two corners of the approximately rectangular lamellipodium (**Figure 2B,** *orange line*), which appears to lie on an axis approximately perpendicular to direction that the cell was traveling recently (*dashed orange line*). The other axis is defined by the rear of the cell through the cell body (*green line*) and is observed to be oriented perpendicular to the direction that the cell is currently traveling (*dashed blue line*). The characteristic asymmetric shape of turning cells was confirmed by creating an average shape from 22 cells selected for their persistent turning behavior, which shares the major features noted in the example shown (**Figure 2B, inset**). Although a previous statistical analysis of shape had shown that only ~1% of total shape variation arises from left-right asymmetry (Keren et al., 2008), within the population of turning cells we found that this particular mode of shape variation was strongly correlated with angular speed (**Figure 2E, Figure S2A**).

Imaging the filamentous actin cytoskeleton driving cell motility in a persistently turning cell (**Figure 2C**) revealed consistent asymmetries in the cytoskeleton that correlated with the observed asymmetries in shape. Our initial expectation was that protrusion would be fastest in the direction the cell was traveling towards, that is on the inside of the turn (Mogilner and Rubinstein, 2010). In this situation actin filament density, *D*, would be expected to be highest on the inner part of the turn as the cell lays down an asymmetrically protruding network. However, we observed instead that the distribution of filamentous actin at the leading edge showed higher F-actin density on the outer wing of the lamellipodium of turning cells (**Figure 2D**), matching the higher aspect ratio on the outer side of the cell. Furthermore, turning cells exhibited significantly faster rates of actin network disassembly in the outer wing of the lamellipodium (**Figure S2B-D**). Thus instead of protruding into turning behavior, cells are actually pivoting around turns with faster lamellipodial protrusion on the outer side of the turn. Quantitatively we note that there is an ~20% increase in F-actin density on the outer side of the turning cell (**Figure 2D**), which can at most account for a few tens of percent increase in the protrusion rate (Keren et al., 2008). Yet, given that the typical radius of curvature of the centroid motion for a persistently turning cell is ~25 *μ*m with a typical cell width of ~40 *μ*m, there must be a ~9 fold gradient in effective net speed from the inside of the cell to the outside. Therefore protrusion asymmetry alone can only be a minor factor in cell turning.

### Turning cells have a stereotyped asymmetric myosin II organization that is necessary to maintain persistent turning

An alternative hypothesis is that whole-cell turning is not primarily driven by asymmetry in actin filament polymerization at the cell leading edge, but rather by asymmetry in the myosin-II driven centripetal flow of actin filaments toward the cell body at the cell rear. We found that the distribution of non-muscle myosin II regulatory light chain at the cell rear, normally found in two similar spots on either side of the cell body in a straight-moving cell (Wilson et al., 2010), is strongly asymmetric in a persistently turning cell, with greater myosin II density on the outer wing of the cell (**Figure 3A, B, *Movie S2***). In addition, the degree of left-right asymmetry in myosin II heavy chain as determined by immunofluorescence among a population of individual cells correlated with a smaller radius of path curvature (tighter turning) for each cell (**Figure 3C**). We also generated a spatial map of myosin activity by comparison of extracted actin networks of turning cells before and after addition of ATP to trigger myosin II network disassembly activity (Wilson et al., 2010), which revealed a pattern of activity that extended more prominently into the outer wing for turning cells (**Figure S2E**), consistent with the faster rate of actin network disassembly observed in this region (**Figure S2D**).

**Figure 3.**
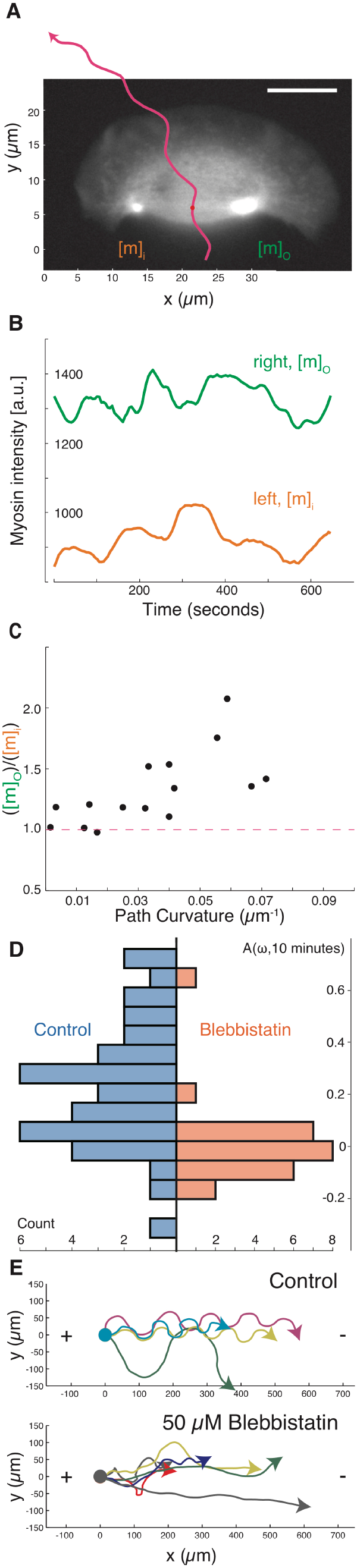
Asymmetric myosin activity drives persistent cell turning. **(A)** A sample image of myosin regulatory light chain-YFP distribution in a turning cell with a counter-clockwise path (*magenta line*). Note that the myosin II density is higher on the outer side of the turning cell. **(B)** Time series of the myosin density in the outer wing, *mo* (*green*), and inner wing, *mi* (*orange*), of the cell from panel A, showing consistently higher myosin on the outer side of this turning cell over time. Note that the vertical axis does not start at 0. **(C)** The relative concentration of myosin II heavy chain as determined by immunofluorescence on the outer and inner sides of cells that were imaged live prior to fixation (vertical axis), plotted against the cell’s path curvature prior to fixation (horizontal axis). Note that cells with a greater degree of turning had an increase in the asymmetry of myosin II localization, correlation coefficient is 0.67 for 15 cells. **(D)** The distribution of calculated auto-correlation of angular velocity with a lag of 10 minutes was calculated for control cells (left/blue) and cells treated with the myosin-II inhibitor blebbistatin (right/orange). Inhibition of myosin-II drastically reduced the number of cells exhibiting persistent turns. **(E)** The trajectories of cells exposed to an electric field of 5 V/cm under control conditions (*top*) and with inhibition of myosin II (*bottom*). All cells migrate towards the cathode on the right, but only cells under control conditions have a periodic overshoot of a straight trajectory suggestive of persistence of a previous angular velocity.

Global pharmacological inhibition of myosin II activity in keratocytes using the small molecule blebbistatin (Straight et al., 2003) decreased the amount of time cells exhibited persistent turning, producing cellular trajectories that were noticeably straighter over long (10 minute) time scales than those of untreated cells (**Figure 3D**). This selective loss of persistent turning in response to myosin II inhibition was also observed in the directed migration of keratocytes toward the cathode in a DC electric field (Allen et al., 2013). Typical cells will turn toward the cathode and then continue to turn persistently, overshooting the straight path predicted by the directional cue of the electric field and therefore producing characteristic strongly periodic oscillations (**Figure 3E, top**). Addition of the myosin II inhibitor blebbistatin inhibits this persistent turning, and therefore cells follow relatively straight trajectories toward the cathode (**Figure 3E, bottom)**. Direct observation of cells turning in an electric field following inhibition of myosin II contractility showed that turning under these conditions was driven morphologically by asymmetric actin polymerization (**Figure S2F**), distinct from the persistent turning behavior described above for unperturbed cells. Thus, pharmacological inhibition of myosin II activity does not prevent the detection of electric fields by keratocytes, but does prevent them from entering into the persistent turning state either spontaneously or following exposure to an electric field.

### Asymmetric myosin II activity drives asymmetric inward actin flow at the cell rear, producing cell turning

Myosin II acts at the rear of the motile cell in part to contract the filamentous actin cytoskeleton, generating net flow of the actin network. Asymmetric actin flow due to the asymmetric myosin II localization described above might lead to cell turning by at least two distinct mechanisms. First, enhanced myosin II activity at the rear at the outside edge of the cell might generate faster retrograde flow relative to the substrate on that side, resulting in actin network slippage relative to the substrate and therefore slower net forward protrusion. Alternatively, enhanced myosin II activity at the outside edge might cause faster inward (centripetal) flow at the rear of the cell only, without affecting retrograde actin flow at the leading edge, such that cells are able to turn by pivoting the orientation of their lamellipodia around their cell bodies. To determine the effects on actin network motion of the persistent asymmetry in myosin II distribution associated with persistent cell turning, we directly measured F-actin network flow using fluorescent speckle microscopy (Danuser and Waterman-Storer, 2006). In the moving cell frame of reference, we found that actin network flow relative to the leading edge was strongly tilted toward the outside rear corner of the cell (**Figure 4A*, Movie* S3**), in contrast to the typical flow pattern for the cell frame of reference in cells moving straight, where the network flows straight backward from the leading edge to the cell rear (Wilson et al., 2010). In the laboratory frame of reference, the F-actin network remained essentially static relative to the substrate at the leading edge, with measurable flow only at the rear of the cell, about 5-fold faster inward on the outside of the turning cell relative to the inside (**Figure 4B**). These observations are consistent with previous observations for keratocyte motility suggesting that newly polymerized actin is tightly coupled to the substrate by integrin-mediated adhesions, resulting in very little retrograde flow at the leading edge (Theriot and Mitchison, 1991; Wilson et al., 2010). Furthermore, this observation rules out the possibility that a hidden asymmetry in relative protrusion is occurring at the leading edge secondary to asymmetric retrograde flow at the leading edge. Instead these results indicate that the lamellipodium of a turning cell can best be thought of as a filamentous actin network that is laid down roughly symmetrically at the front but acted on asymmetrically at the rear by inward myosin II-dependent contractility to produce large-scale cellular turning.

**Figure 4.**
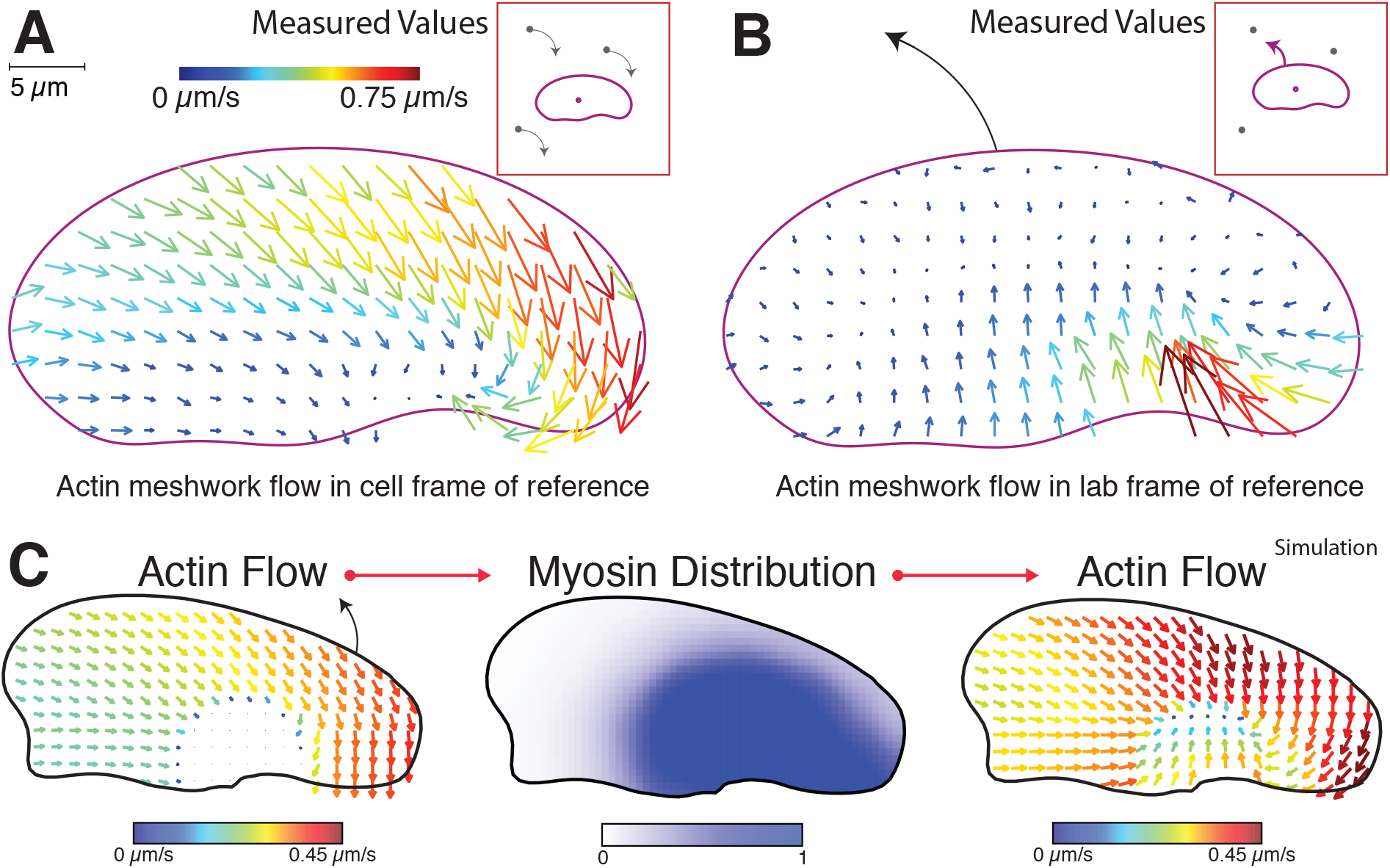
Asymmetric centripetal actin network flow in turning cells produces an asymmetric myosin distribution. Experimentally determined vector maps representing fluorescence speckle microscopy measurements of the flow of the F-actin network in the cell frame of reference **(A)** and in the lab or substrate frame of reference **(B)** for a cell turning counter-clockwise. Results are representative of measurements made in 12 separate cells. Graphical depictions of each frame of reference are presented in insets. Scale bar indicates 5 *μ*m, and vector arrow size and color scales with magnitude of flow speed. **(C)** From the depicted assumed asymmetric shape and asymmetric flow of the actin network in the cell frame of reference (left), the distribution of myosin inside the cell was generated (center). Then using the assumed asymmetric shape and this myosin distribution, the kinematic flow of actin in the cell frame of reference was recalculated (right). In this model the asymmetric flow of actin toward the outer side of a turning cell is sufficient to bias myosin to the outer side, which in turn is sufficient to approximately recreate the original asymmetric flow of actin. Scale bars below each plot indicate vector magnitude strength and local myosin density.

To determine if these mechanical asymmetries could be sufficient to reproduce turning behavior in keratocytes, we employed a simulation of a mechanical model of the keratocyte lamellipodium as previously tested and calibrated for straight steady-state motility (Barnhart et al., 2011) and for the process of polarization and motility initiation (Barnhart et al., 2015). In brief, this model uses the balance of three essential forces (myosin II contraction, adhesive drag and actin shear) to determine the cell’s mechanical behavior. Cell shape is set by the balance between the rate of actin polymerization-driven protrusion normal to the cell boundary, *V_p_*, which is limited by membrane tension, and myosin II-driven retraction of the network normal to the cell boundary, *U_⊥_*. In this model, myosin II acts to generate contractile stress applied to a viscous actin network with viscous resistance to actin flow created by adhesion to the substrate. For simplicity, in the model the resistance of adhesion to the flow of actin (*U*) is linearly proportional to the local adhesion strength, *ζ*, and myosin contraction is linearly related to the local density of myosin II, *M*, which is established by the transport of myosin bound to the F-actin network traveling towards the cell rear in the frame of reference of the cell (Svitkina et al., 1997).

To model the effect of asymmetries on the motile cell, we first used a simplified fixed boundary model (**Figure S4J**) and considered a steadily turning cell with a fixed shape derived from our observations of a stereotyped turning cell (**Figure 2**) and an assumption of spatially symmetric adhesive drag (**Figure S3A**). From the point of view of the turning cell, we had experimentally observed that the actin network slides diagonally inward on the inner side of a turning cell and straight back on the outer part of the cell (**Figure 4A**). Starting with the initial condition of asymmetric actin flow matching the experimental observation, we found that this was sufficient to polarize the distribution of myosin to the outer edge at steady state (**Figure 4C**). Moreover, when we instead used this simulated asymmetric distribution of myosin as the initial condition within the fixed boundary model, we generated steady-state actin flow fields that matched the original experimental observations (**Figure 4C, right)**. These simulations show that, as a cell turns, the localization of myosin II can become biased to the outside due to its coupled transport with the flow of the actin network in the cell frame of reference. At the same time, the asymmetric myosin II localization can then contribute to a positive feedback loop, reinforcing the asymmetry of the actin network flow. This kind of positive feedback loop is consistent with our behavioral observations on the ability of these cells to enter into a persistent turning state.

### Traction force asymmetry implicates asymmetric cell adhesion during turning

Thus, we find that actin network flow and myosin contractility alone can establish a positive feedback loop reinforcing left-right asymmetry in turning keratocytes, but it is also known that cell-substrate adhesion in migrating cells is coupled to both actin network flow and to myosin II contractility (Gardel et al., 2010). We therefore wished to explore whether asymmetries in adhesion might also contribute to the persistent turning state. To this end, we used our force-balance model using a fixed asymmetric cell shape, as described above, to simulate cell turning assuming either a spatially constant adhesion distribution as previously discussed or an alternative distribution with weaker adhesions on the outside of the turning cell. We found that between these two adhesion strength distributions there was little change in the resultant simulated distribution of myosin II or actin network flow (**Figure S3B**). However, simulated traction forces were strongly dependent on the assumed adhesion distribution. The simulation with a constant adhesion distribution found high traction forces on the outside of the turning cell, with weaker traction on the inside (**Figure 5A),** while the asymmetric adhesion distribution simulation found traction forces that were weaker on the outer side despite greater speed of centripetal actin flow (**Figure 5B**).

**Figure 5.**
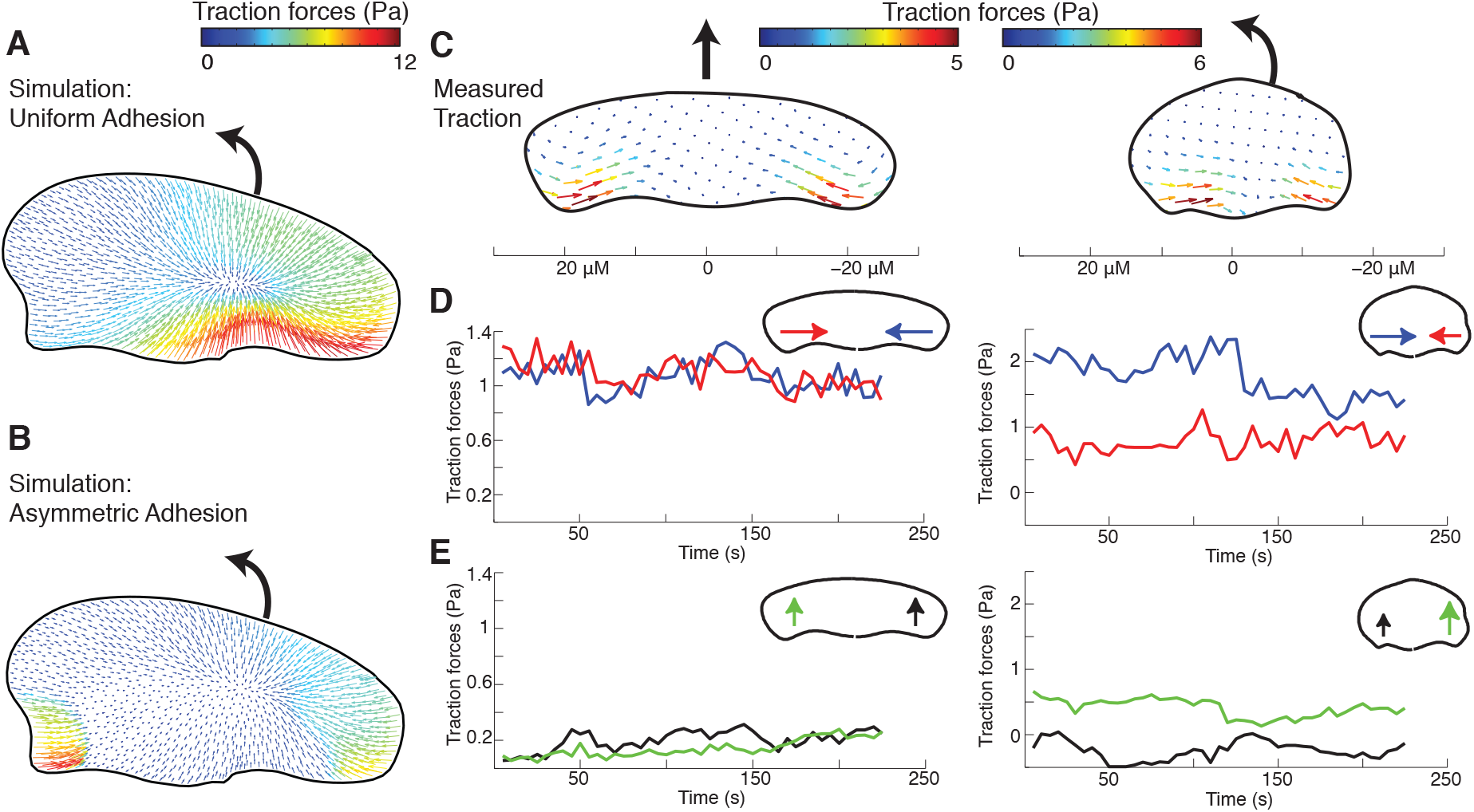
Turning cells apply asymmetric traction forces. Vector maps of the simulated traction forces exerted on the substrate from a cell turning clockwise with an asymmetric shape and an initial asymmetric flow of the F-actin network as depicted in Figure 4C using either a symmetric distribution of adhesion strength **(A)** or an assumption of higher adhesion strength on the inner side of the turning cell **(B)**. Full simulation results are available in Figure S3B. Vector color and size is set by magnitude of traction force as depicted in scale bar. **(C)** Vector maps of average experimentally measured traction forces created by a cell migrating along a straight path (left) and a cell turning counter-clockwise (right). Scale bar presented below is in microns. **(D)** Time series of the spatial average of the inward component of traction forces on the left (*red*) and right (*blue*) sides of the same cells as in panel C. Note that turning cells have persistently higher inward traction forces on the inside of the turn when compared to the outside, which matches simulations performed with higher adhesion strength on the inside of a turning cell. **(E)** Time series of the spatial average of the forward component of traction forces on the left (*green*) and right (*black*) sides of the same cells. Note that turning cells have persistently higher forward traction forces on the outer side of the turning cell when compared to the inner side, again matching simulations with left-right adhesion asymmetry.

With these predictions in hand, we measured the traction forces that a turning cell applied to the substrate through its actin network using traction force microscopy (Sabass et al., 2008) in order to experimentally distinguish between these two possibilities (**Figure 5C**). We found that motile keratocytes primarily generated traction forces orthogonal to the direction of motion, confirming prior results (Oliver et al., 1999). Cells that moved straight showed midline symmetry in the inward forces exerted on the substrate (*T_⊥_(in/out)* =1.1±0.3, n = 4) whereas turning cells had an increase in inward-directed traction forces on the inner side (*T_⊥_(in/out)* =1.7±0.7, n = 5), matching the predictions of the simulation with the assumption of asymmetric adhesion (**Figure 5C-D**). In addition, forces in both the simulation and experimental measurements were found to have a more forward orientation at the outside rear for turning cells, as compared to either the inside rear of a turning cell or the rear of a cell moving straight (**Figure 5E**). The net force on the cell on the surface under both conditions was approximately zero, as expected. Mechanistically, the observed large inward traction force on the inner side of the turning cell must result from the gripping traction from high local adhesion at the pivoting point, which is balanced by opposing traction throughout the rest of the cell. The forward component of the traction force at the rear of the outer side of the cell is a “slipping” resistive adhesion force due to forward locomotion, and is balanced by the “gripping” retrograde traction force spread over the wide leading edge.

Together, our observations of faster inward actin network flow on the outside of a turning cell (**Figure 4B**) coupled with stronger traction force on the inside of the turning cell (**Figure 5C-D**) are consistent with a non-linear “stick-slip” mechanism for traction force generation (Sabass and Schwarz, 2010), where traction stresses are proportional to F-actin flow at low F-actin flow speeds and inversely proportional to flow at high speeds, as has been experimentally observed in mammalian epithelial cells (Gardel et al., 2008) as well as in keratocytes undergoing spontaneous symmetry-breaking during movement initiation (Barnhart et al., 2015). As the inward flow of the actin network at the keratocyte rear is driven by myosin II contractility (Wilson et al., 2010), this sets up a second feedback loop, where increasing the myosin contraction of the viscous actin network will at first act to increase the strength of adhesion until a critical point where contraction and F-actin flow speed are too high to be sustained, at which point increasing contractility weakens or breaks adhesions, further facilitating rapid flow of the actin network on the outer side of the cell.

### Local modification of adhesion or myosin contractility is sufficient to induce cell turning

From these observations, we predicted that experimental perturbation of either local myosin contractility or local adhesion on one side of the rear of a moving keratocyte should be sufficient to induce cell turning. Indeed, we found that direct local application of the myosin II activating serine-threonine phosphatase inhibitor calyculin A using a micro-needle to one side of individual cells was sufficient to induce cells to turn away from the site of drug application (*dθ* = 46 ± 24 degrees, **Figure 6A**), unlike control dye administration (*dθ* = −10 ± 50 degrees). Similarly, when we examined the migration of individual cells crossing boundaries between substrates of normal to low adhesivity (Barnhart et al., 2011) we found that asymmetry in adhesion could cause cell turning towards the higher adhesion substrate (**Figure 6B,C, Movie S4**). The average induced turn was ~90 degrees and as expected had an apparent dependence on the angle of incidence, which dictates the degree of adhesion asymmetry. Cells crossing the boundary in the opposite direction (from low to normal adhesivity) also turned toward the substrate of higher adhesivity (**Figure 6C**). Importantly, we found that directionality changes predominantly (>70%) occurred as the rear of the cell crossed the adhesion boundary. This confirms that asymmetric myosin II contraction of the actin network at the cell rear alone or asymmetric coupling of the myosin II mediated centripetal actin flow to the substrate are both sufficient to trigger a cell to transiently turn away from the side of faster inward actin network flow. However, neither treatment could routinely induce a persistent turning state where cells generated circular trajectories as described above; instead most cells returned to a fairly straight path after transient perturbation (**Figure 6A,C**).

**Figure 6.**
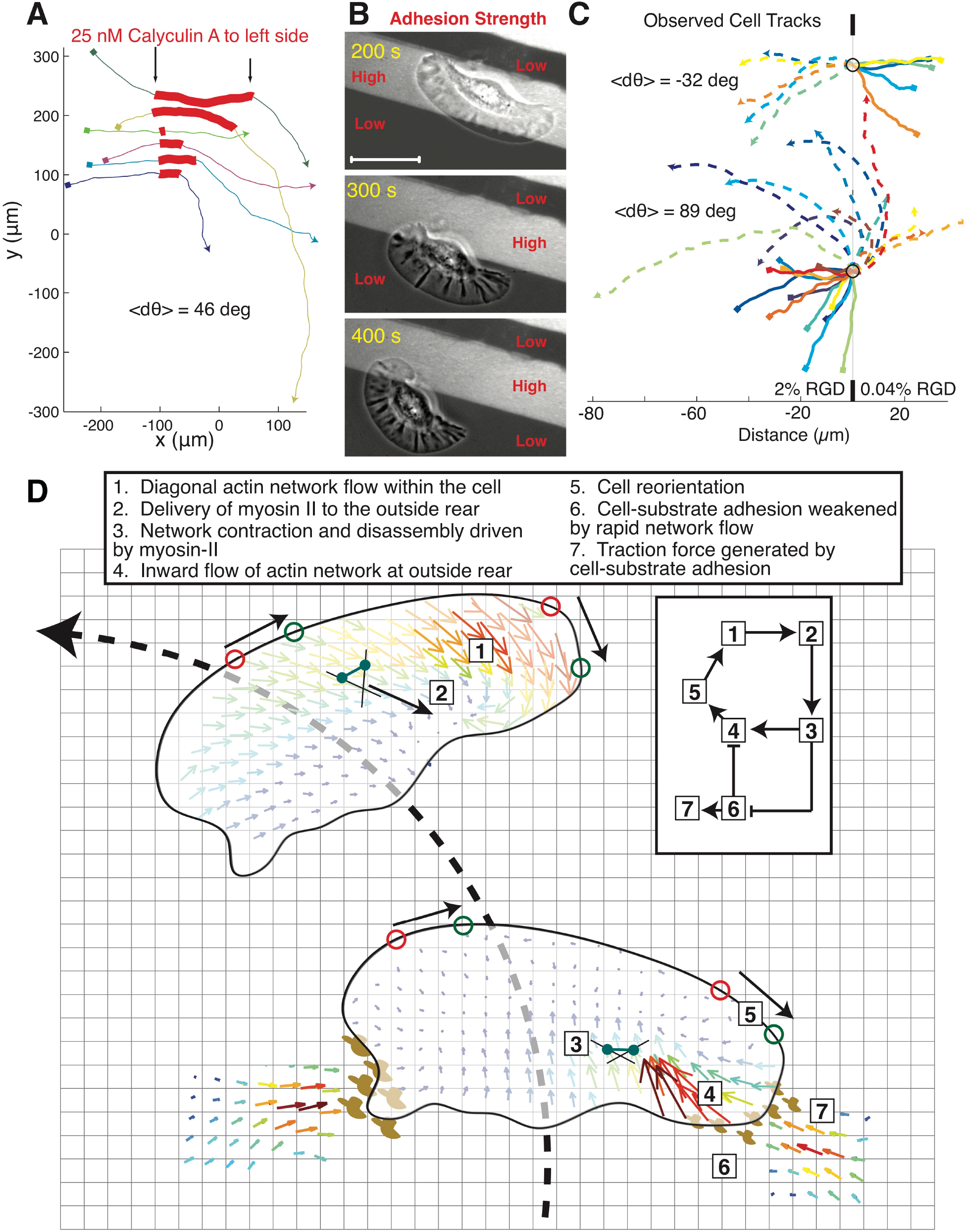
Asymmetries in myosin II activity or cell-substrate adhesion are sufficient to induce turns. **(A)** The trajectories of a set of cells that were asymmetrically exposed to calyculin locally on the left side of the cell during the portion of the trajectory marked in red. Local calyculin exposure induced cells to turn away from the side of upregulated myosin II activity. Trajectories start at squares and proceed from left to right. Each color indicates a different cell. **(B)** Images of a single cell crossing a boundary between a 2% RGD normal adhesion substrate (*light*) on to a 0.04% RGD low adhesion substrate (*dark*), causing the cell to turn toward the high adhesion side of the cell when adhesion at the rear becomes unbalanced. Scale bar indicates 10 *μ*m. **(C)** Trajectories of a set of cells crossing a boundary of normal adhesion density (2% RGD) to low adhesion density (0.04% RGD) on the bottom and from low adhesion density to normal adhesion density on the top. Start positions are marked with small squares, dashed lines indicate trajectories after hitting boundary. All trajectories are centered on the boundary collision point marked with circle. Cells either reflect off the adhesion boundary or refract towards the side of higher adhesion, where the change in direction is dependent on the incident angle with the boundary. Scale bar on bottom indicates distance in microns. **(D)** A model of cell turning in keratocytes. In the turning cell the diagonal outward facing direction of actin flow *[1]* increases the delivery of myosin to the outer side of the cell *[2]* increasing myosin contractility *[3]* and the centripetal flow of actin slipping over the substrate on the outer side of the cell *[4]*, producing cell turning which in turns drives the diagonal flow of actin to the outer edge of the cell *[1]*. Increased myosin contractility on the outer side of the turning cell *[3]* also weakens adhesions *[6]* decreasing traction forces on the outer edge of the cell *[7]* and further promoting the centripetal sliding of the outer actin network over the substrate *[4]* and consequently turning.

So far, our experimental work and simulation results are all consistent with a complete mechanical model of cell turning involving two feedback loops: asymmetric actin/myosin II contraction driving asymmetric inward actin flow that is perpetuated both by asymmetric delivery of myosin II and asymmetric engagement of adhesions (**Figure 6D**). However, two questions remain at this point; are these interactions sufficient to determine the characteristic asymmetric shape of turning cells (**Figure 2**)? And, what explains the coexistence of both persistently turning and randomly meandering keratocytes in cell populations (**Figure 1**)?

### A mechanical model can produce realistic cell turning and explains variation in cellular trajectories

We returned to our simulation to explore these remaining questions, adding a freely deformable cell boundary so that we could explicitly examine the evolution of asymmetric cell shape over time. In the free boundary model, the movement of each point along the cell boundary is determined by the balance of actin polymerization-driven protrusion (which in turn is limited by membrane tension) and myosin II-driven contraction, as described in the supplementary discussion (**Figure S4J**). Starting from a stable symmetrical shape, the simulated free boundary cell incorporating only the feedback loop between actin flow and myosin II contractility (but not the second feedback loop connecting contractility to adhesion) evolved into a polarized, motile configuration with a relatively straight, slightly meandering path (**Movie S5**). Adding a fixed and persistent asymmetry in adhesion strength (**Figure 7A**) or myosin distribution (**Movie S6**) was sufficient to induce an asymmetric cell shape, actin flow, and traction pattern, that were qualitatively similar to our experimental observations for persistently turning cells. Examining the simulation across a range of fixed adhesion asymmetries, (*∆ζ*), we found that the cell would develop a steady shape with stable asymmetries in myosin II distribution (*m*), and angular speeds (*ω*), that were approximately linear functions of the adhesion asymmetry (**Figure S4A-C**), *ω≈αΔζ* and *m≈βω*, thus reproducing the experimental observation that the degree of left-right myosin II asymmetry correlated with a smaller radius of curvature (**Figure 3C**).

**Figure 7.**
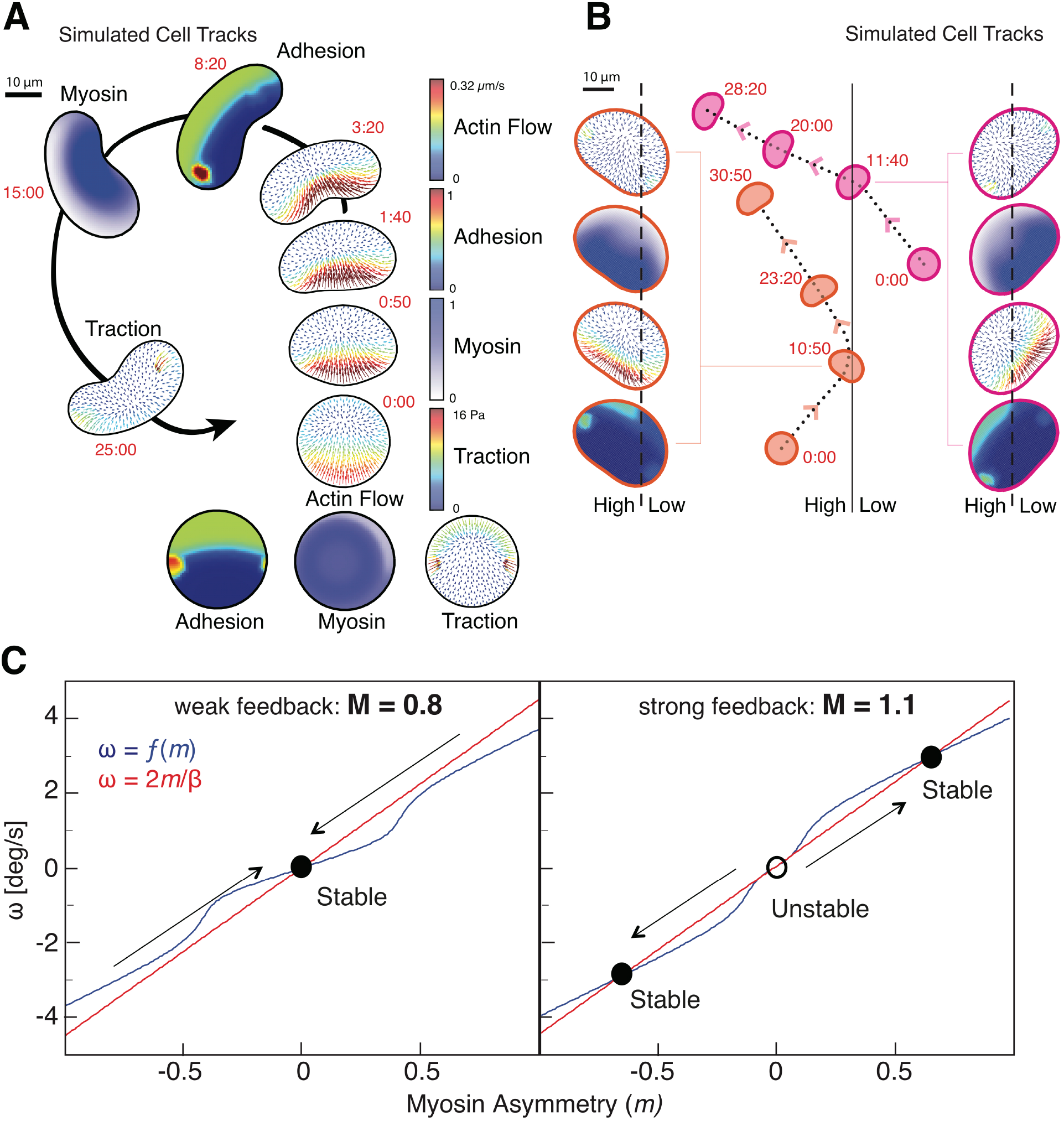
Characteristics of cell turning can be reproduced with a mechanical model. **(A)** The results of a free-boundary simulation of cell migration where cell shape and migration evolved with the initial assumption of increased adhesion strength on the left side of the cell. Cell outlines are shown at time points as labeled in minutes (*red*), with the first two time points not presented in their actual location in space. Note that we observe expected changes in cell shape, actin flow (*3:20*), adhesion asymmetry (*8:20*), myosin distribution (*15:00*) and traction forces (*25:00*). **(B)** Simulation with free-boundary model of cell migration of cells migrating from high to low adhesion (*orange contours*) and from low to high adhesion (*magenta contours*). Cell shape and parameters at the point of crossing are presented on the sides as labeled, with scale bars as in panel A. Matching experimental observations, cells turn towards the side of the substrate with higher adhesion in both cases. Scale bar indicates 10 *μ*m. **(C)** From the combination of the lisnear relationship between angular speed and myosin asymmetry dictated by asymmetric delivery of myosin II to the outer side of the cell inducing turning (*red*) and the non-linear relationship between angular speed and myosin II asymmetry dictated by the weakening of adhesion by myosin II contractility according to a stick-slip model (*blue)*, the stable turning state of a cell can be determined. When myosin II contractility is too weak to break adhesion (M = 0.8) then a single stable solution can be obtained as would be seen for a meandering cell (*left)*. When myosin contractility is stronger (M = 1.1) and weakens adhesions promoting further turning, then a cell will be stable in either of two persistent turning states (*right)*.

Next, we explicitly examined the importance of the stick-slip interaction between local myosin contractility and local adhesion (**Figure S4D**). In simulations where the feedback between contractility and adhesion was weak, transient asymmetries in adhesion alone produced relatively weak turning behavior, whereas strong negative feedback produced turning in persistent circles (**Figure S4E-H**). The full free boundary model including both feedback loops was also strikingly able to reproduce the behaviors we observed for cells crossing boundaries from high to low or low to high substrate adhesivity (**Figure 7B, Movie S7A-B**). In contrast, when we simulated the effect of increasing protrusion asymmetrically on one side of the cell, we found that the cell would deform but not turn (**Movie S8**), consistent with our overall conclusion that persistent keratocyte turning is driven by the mechanics of retraction at the cell rear and not by graded protrusion at the cell front. In addition, we simulated the effect of galvanotaxis by adding the observed influence of the applied electric field on the cell leading edge (**Figure S2F**) to bias the cell’s protrusion direction toward the cathode to the model. Solutions of this model reproduced the observed effect of periodic oscillations of the cell trajectory (**Figure S4I**) The explanation is simple: the cell begins to turn persistently and overshoots the direction preferred by the electric field guidance, but when the cell deviates too far from the cathode direction, the growing pull of the leading edge slows the angular velocity enough to weaken the positive feedback between turning and myosin II asymmetry. The cell starts turning back toward the cathode, myosin II redistributes again, and the restored feedback locks the cell in the turning state again until the overshoot in the other direction becomes too great, and so on. Simulating the effect of weakening myosin II contractility shows that the cell trajectory stops oscillating without this feedback, also in agreement with our experimental data (**Figure 3E**).

Finally we used our simulations to turn to the question of why some individual keratocytes enter into persistent turning states while others do not. In order to generate a sufficiently large number of simulated cellular trajectories to examine their statistical behavior, we used the simple relationships between adhesion asymmetry, myosin II asymmetry and angular velocity (**Figure S4A-C**) as well as the stick-slip relation (**Figure S4D**) to derive a simple system of stochastic equations that are easy to solve (see *Supplementary Material*). This stochastic simulation was able to reproduce multiple features observed in the experimental cell trajectories, most strikingly the puzzling plateau in the mean squared displacement of angular speed over time (**Figure S1D**). We simulated trajectories for cells starting with a wide range of initial myosin II asymmetry states, using various values for the overall strength of myosin II contractility in the cell, and examined their approach to steady-state behavior (**Figure S4E-H**). This analysis predicts that, in the presence of sufficiently strong positive feedback between turning and myosin II asymmetry, and sufficiently strong negative feedback between contraction and adhesion strengths, the cell becomes mechanically bistable (**Figure 7C, right, Movie S9**): it turns persistently either clockwise or counter-clockwise with a particular characteristic angular velocity. This non-intuitive result satisfyingly explains the symmetric bimodal distributions of angular speeds for individual persistently turning cells, where the absolute magnitude of the angular speed was typically very constant as the cell switched from clockwise to counter-clockwise turning behavior (**Figure 1C,E; Figure S1C, top**). In contrast, our model predicts that if the combination of these feedbacks is too weak, the cell stably goes straight, meandering only due to the stochastic noise (**Figure 7C, left**). Thus, cells with weaker overall myosin II contractility are predicted to have meandering trajectories without entering persistent turning states, exactly as we observed experimentally for cells where myosin II was globally inhibited (**Figure 3D**). This result also provides a satisfying explanation as to why local and temporary activation of increased myosin II contractility or increased adhesion on one side of the cell rear could induce single turns but could not drive cells into the persistent turning state (**Figure 6A-C**); these cells, which were moving fairly straight before perturbation, are likely to be in a relatively low contractility state and therefore return to their single stable point after transient perturbation (**Figure 7C**). The suggestion that individual keratocytes in a population may have slightly different levels of myosin II activity, driving them into different net behavioral states with respect to persistent turning, is highly consistent with our previous observations of cell-to-cell variation in VASP activity (Lacayo et al., 2007) and total F-actin (Keren et al., 2008) driving large-scale variations in lamellipodial morphology and cell speed.

## Discussion

From this combination of experimental observations of persistently turning keratocytes and computational simulations we can now derive a complete mechanical model for how these cells turn (**Figure 6D**). Turning starts with asymmetric actin flow created by asymmetry in adhesion and/or asymmetry in myosin II contractile activity at the cell rear. This rotates the “corners” that define the lamellipodial leading edge, which effectively rotates the protruding actin network relative to the cell body and changes the direction of movement of the front of the cell as a result of mechanical changes at the rear. The resulting turning causes the filamentous actin network and the attached myosin II to sweep preferentially into the outer wing, which maintains the flow asymmetry, enhancing further myosin II accumulation in the outer wing, and producing positive feedback. This type of kinematic feedback loop of an inhibitory molecule from the front to the rear of the turning cell has previously been predicted in amoeboid cells such as *Dictyostelium discoideum* to produce random turns that become uncorrelated over long time scales (Nishimura et al., 2012). Similarly we found that the positive feedback between the kinematics of turning, myosin II redistribution and actin flow asymmetry are not strong enough by themselves to be self-sustaining, unless the increased myosin II contractility is sufficient to weaken or break adhesions on the outer side of the cell, creating a more locked-in asymmetry in both myosin contractility and adhesion strength.

We also experimentally observed a peaked F-actin distribution on the outer side of the turning cell leading edge, which typically signifies a faster protrusion rate. This is a likely result of the mechanics of cell turning, as any given molecule limiting actin polymerization that diffuses along the leading edge will be biased to outer side of the turning cell by rotational drift (Lacayo et al., 2007). Increased myosin II activity and concomitant F-actin disassembly (Wilson et al., 2010) will also produce a local increase in monomeric G actin on the outer side of the turning cell, further promoting faster protrusion on the outer side. However, quantitatively these changes in protrusion are not sufficient to explain the degree of turning seen experimentally, and using our computational free-boundary simulations we were able to reproduce asymmetric cell shape and curved trajectories without an asymmetric distribution of either F- or G-actin (**Figure 7A**). Thus, these asymmetries in protrusion at the leading edge appear to play only a minor role in cell turning, becoming dominant only when the role of myosin II contractility is inhibited. It is interesting to note that local photoactivation of G-actin sequestering thymosin *β*-4 was reported to induce keratocytes to turn towards the side of less protrusion, though without directional persistence and with a reported decrease in contractility on the thymosin *β*-4 exposed side of the cell (Roy et al., 2001). In confirmation, our model simulations indicate that, with symmetric contractility or adhesion, asymmetric protrusion alone deforms the shape of the cell but produces limited turning.

Previous attempts to understand the mechanics of keratocyte turning behavior (Oliver et al., 1999) focused on the asymmetry of traction forces, insightfully decomposing the forces into propulsive, pinching and resistive parts. However, this previous study interpreted the traction asymmetry from the point of view of an effective torque rotating a rigid cell body, and did not address the cytoskeletal asymmetries underlying the turning mechanism. Our analysis suggests that the traction forces are coupled to the asymmetric viscous contractile actin-myosin network within the free boundary of the cell, and the key to cell turning is the combination of the force asymmetries with the kinematics of actin flow and cell shape deformations. It is not the case that the turning cell rotates as a rigid body; instead the asymmetrically dynamic lamellipodium slides laterally relative to the cell body to produce a change in the net direction of whole-cell movement.

Our work identifies how the mechanical actors of cell migration work together to follow a cell’s internal compass in generating left-right asymmetry while still maintaining constant front-rear polarization. In vivo, these cells will be under the constant influence of external tactic cues. In neutrophils and *Dictyostelium discoideum*, these tactic cues have been thought to act to promote migration by co-opting the internal compass at the front of the cell to promote protrusion through secondary chemical messengers (local excitation) and to suppress migration at the cell rear by activation of myosin II based contraction (global inhibition) (Gutierrez et al., 2011; Xiong et al., 2010). However, we have found that contractility at the rear of the cell may also act asymmetrically to direct the motion of the front, due to the internal reorientation of the lamellipodium, i.e. rear-wheel steering. Because our analysis has focused primarily on cells turning persistently at steady state, we are not able to determine which events are most likely to initiate the cascade of positive feedback loops we have described here, though it is clear that it is possible to trigger at least some part of the rear-wheel steering process by altering either contractility or adhesion asymmetrically at the cell rear. In a parallel project in human neutrophils (Tsai et al., submitted), we have found that asymmetrical delivery via actin network flow of myosin II to the outside rear of turning neutrophils follows patterns very similar to those described here in keratocytes, with the interesting distinction that the left-right myosin II asymmetry at the cell rear is not persistently maintained, so neutrophils do not enter into persistent turning states. For neutrophils, the first step in both spontaneous and induced cell turning appears to be reorientation of the actin network flow at the leading edge, leading to asymmetric myosin II delivery and the same sequelae as described here for keratocytes.

Several recent lines of experiment have implicated large-scale actin network flow from the front to the rear as a conserved property of many migrating cell types that can serve to integrate cell-scale behavior (Callan-Jones and Voituriez, 2016). In particular, actin cortical flow driven by myosin II contractility in amoeboid cells in the developing zebrafish embryo is sufficient to generate robust self-reinforcing front-rear polarity (Ruprecht et al., 2015). Across a wide variety of motile cell types, there is a strong correlation between the speed of cell movement and directional persistence, which can be simply modeled with the proposition that the net rearward movement of the flowing actin network transports molecules responsible for the reinforcement of front-rear polarity to the back of the moving cell (Maiuri et al., 2015). Our present findings on persistent turning in keratocytes reveal similar mechanisms at play in motility driven by lamellipodia, where the primary factor transported by the actin network to determine front-rear polarity is simply myosin II itself (Ofer et al., 2011; Svitkina et al., 1997; Wilson et al., 2010; Yam et al., 2007). The implication that left-right asymmetries in network flow and therefore in delivery or activation of myosin II might also contribute to cell turning in other motile cell types beyond keratocytes and neutrophils remains to be explored.

## Conclusions

Here we have presented an analysis of the asymmetries that are utilized by the motile keratocyte to change direction in space. Our experiments demonstrate that directionality is dictated primarily at the rear of the cell through a mechanical mechanism featuring two feedback loops between actin network flow, myosin II contractility and adhesion to the substrate. We found that a mechanical model incorporating these features can reproduce actual cellular shapes and trajectories in space. For these cells, asymmetries at the protruding leading edge play a surprisingly minor role in determining the direction and magnitude of spontaneous turning – instead, cells steer from the rear.

## Acknowledgements

We thank Zachary Pincus, Washington University School of Medicine, for development of the Cell Tool image analysis software, Ulrich Schwarz and Benedikt Sabass for the development of computational methods to calculate traction force, and Aaron Straight for the Xenopus regulatory myosin light chain YFP fusion plasmid. This work was supported by the National Institutes of Health and the Howard Hughes Medical Institute.

